# High Precision Quantification of small RNA Slicing Activity - Native Index Ligation-based Targeted Degradome Sequencing (NIL-TDS)

**DOI:** 10.1101/2025.09.30.679503

**Authors:** Bernhard T. Werner, Sabrine Nasfi, M. Lienhard Schmitz, Manar Makhoul, Jens Steinbrenner, Patrick Schäfer

## Abstract

RNA interference (RNAi) is an effective and precise regulatory mechanism in eukaryotes in which small RNAs mediate endonucleolytic slicing of complementary target mRNAs. Despite the potential of RNAi for human therapeutics and crop bio-protection, analytical platforms to quantitatively validate the slicing activities of small RNAs remain limited. Here, we present NIL-TDS, a cost-effective method that combines RNA ligase-mediated PCR with Nanopore Sequencing for direct, rapid, and high-resolution detection and quantification of sRNA-mediated slicing events. Using NIL-TDS, we quantitatively detected minute changes in *Ath*-mir400 mediated slicing of *PPR1* in heat and salt stressed Arabidopsis plants. We further demonstrate the broader applicability of NIL-TDS by detecting rare slicing events in a mammalian system. Of relevance for malignancy of certain cancers, and tumor progression and metastasis, NIL-TDS further confirmed *miR-196-HOXB8* interactions in lung cancer cells and discovered a novel miR-7162-HOXA10-AS slicing site. These findings demonstrate the sensitivity of NIL-TDS for uncovering small RNA mediated regulatory mechanisms of gene expression and disease progression in eukaryotes.

**GRAPHICAL ABSTRACT:** 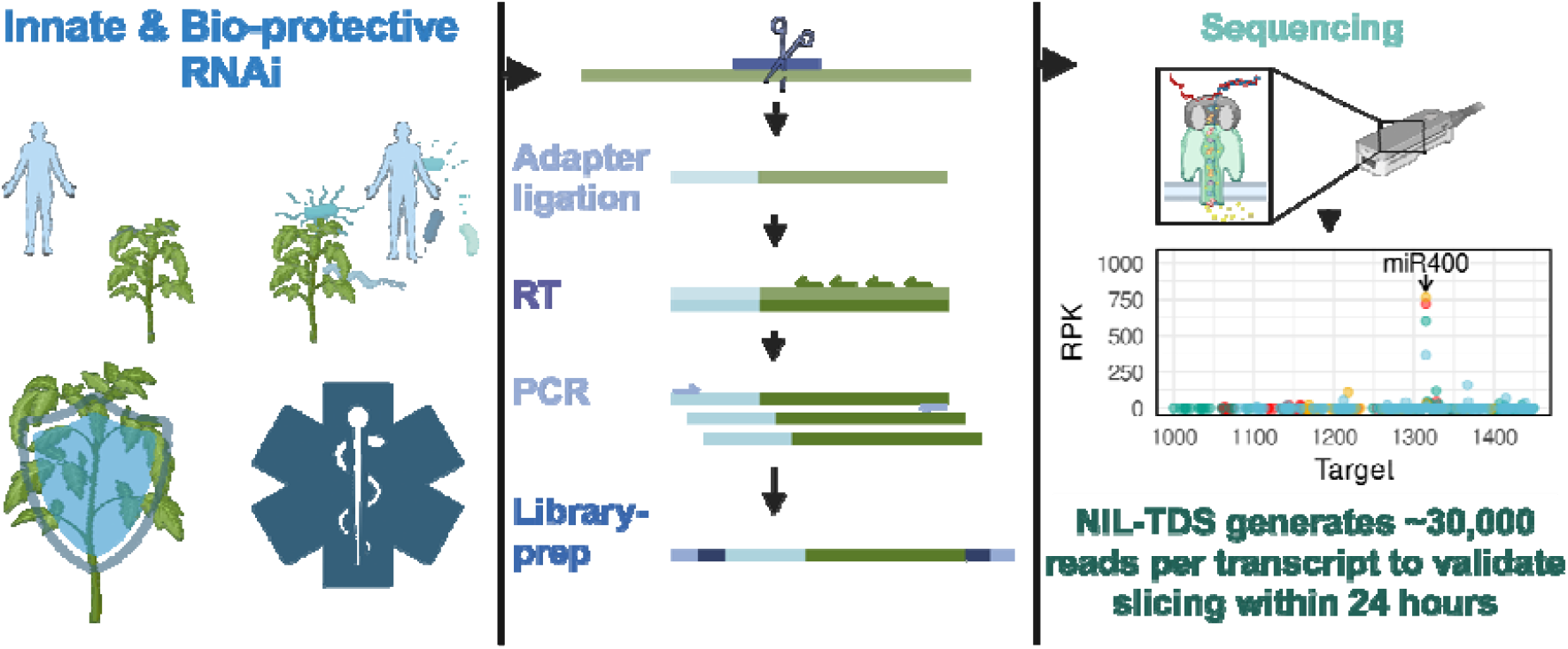

## INTRODUCTION

RNA interference (RNAi) is a conserved mechanism in eukaryotes in which small regulatory RNAs (sRNA) guide the RNA-induced silencing complex (RISC) to slice and thus degrade complementary mRNAs (Shabalina and Koonin 2008). sRNAs are short (∼18-30 nt), double-stranded RNAs that regulate mRNA stability based on sequence complementarity to control fundamental biological processes in eukaryotes. These functions range from growth, development, stress adaptation, and immunity, to cellular processes such as genome stability (Ketting 2011). Variations in this sequence-specific functionality that affect sRNA-mRNA complementarity or accessibility can lead to diseases and developmental defects in animals and plants. For example, a target site mutation in the B-cell activating factor gene, essential for the survival and function of B lymphocytes (a type of white blood cells), abolishes RNAi-mediated suppression, which correlates with an increased risk of autoimmune diseases (Steri et al. 2017). Similarly, a synonymous polymorphism in the target site of microRNA-196 (miR-196) in the immunity-related GTPase M is associated with Crohn’s disease (Brest et al. 2011). In plants, miR164 suppresses the initiation of axillary shoot buds, and mutations in this miR lead to significant morphological changes (Raman et al. 2008).

Designing and engineering sRNAs for RNAi application allows to specifically regulate gene levels with the unique potential to develop customised bio-protective and therapeutic agents for plant and human health (Cai et al. 2018) (Fig. 1A). However, this potential must account for the complexity of sRNA-mRNA target interactions. Depending on the degree of sRNA complementarity, the mRNA target is degraded or translationally inhibited by a process that is still not fully understood (Kern et al. 2019). In addition to mRNA target complementarity, abundance, as well as sRNA subcellular localisation, and interactions with other RNAs or RNA binding proteins (e.g. sponging effect) shape sRNA-mRNA interactions (Diener et al. 2023). Considering this complexity and the continuous progress in next generation sequencing-based discovery of sRNA candidates (Benesova et al. 2021), the development of rapid and highly sensitive methods for the quantification of sRNA-mediated slicing activities is urgently needed. Currently RNA ligase-mediated rapid amplification of cDNA ends (RLM-RACE) (Fig. 1B-C) is the standard procedure for the detection and validation of slicing-based RNAi activity of candidate sRNAs. It has, however, some significant shortcomings. In RLM-RACE, an RNA adapter is ligated to the 5’-monophosphate of the sliced RNA, and this adapter is used for reverse transcription Liave et al. (2002), German et al. 2008). Subsequently, gene-specific primers amplify cDNA ends in a nested PCR reaction. The resulting products are then cloned into a vector, followed by DNA sequencing (Liave et al. 2002). Each clone represents only one RNA-end read, and because of the costly and laborious nature of the procedure, only a few reads are commonly generated for a given gene. As a results, RLM-RACE is a low-throughput approach for validating known candidates that is prone to PCR amplification biases, and does not provide quantitative, statistically confirmable insights into cleavage efficiencies. High-throughput sequencing approaches, such as parallel analysis of RNA ends (PARE) (Zhai et al. 2013) are also used for detection and validation of sRNA slicing activity, where a type II restriction enzyme produces 20-nucleotide long tags, which are further ligated to another 3’-adapter for a second PCR prior to sequencing. Although PARE can potentially provide data on the entire degradome, the detection of slicing events correlates with transcript copy numbers, resulting in insufficient coverage for most, especially low abundant, transcripts (German et al. 2008. Currently, a standard Illumina sequencer can produce billions of reads in the span of a single day, making it compatible for PARE (Wetterstrand 2024; Kumar et al. 2024). However, this technique is not cost-effective for validating single target genes, which are often required for the development of RNAi-based therapeutics. The lack of read depth either in RLM-RACE or in PARE generally results in qualitative data and prevents statistical validation of sRNA slicing events making both methods limited in detecting rare cleavage products and potentially leading to an underestimation of sRNA-mediated slicing events. Fortunately, advancements in sequencing technologies supports the development of alternative approaches (Kumar et al. 2024). The ONT Flongle sequencing platform is an affordable hardware for direct DNA sequencing, which significantly reduces library preparation time. Moreover, throughput and run time are scalable, and single runs to analyse slicing of selected candidates can be operated at competitive costs (De Cesare et al. 2024).

**Figure 1:**
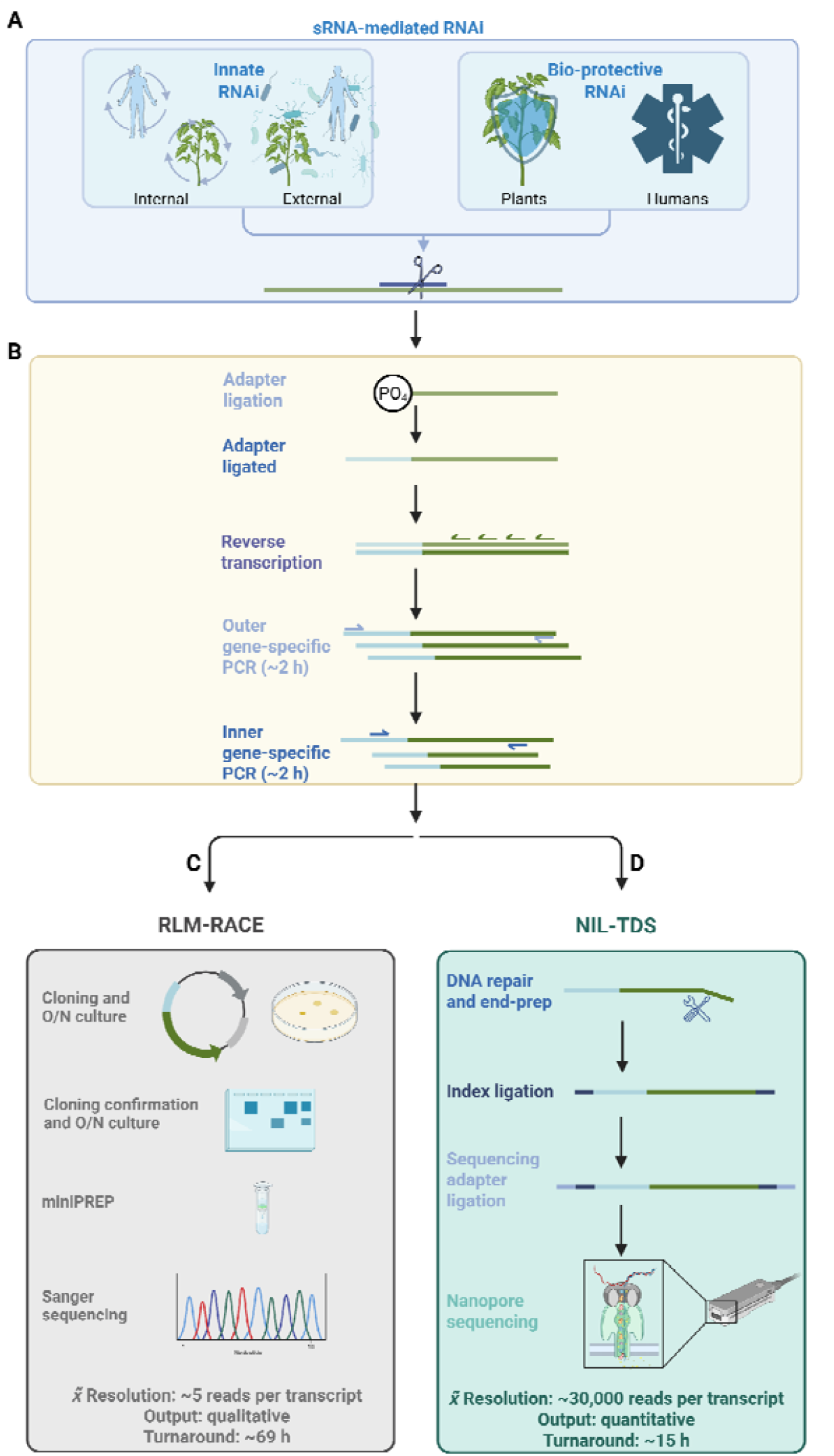
Workflow of quantitative NIL-TDS in comparison to qualitative RLM-RACE. sRNAs trigger RNAi in eukaryotes and can protect plant and human health (A). RNAi generates characteristic 5’-monophosphate RNA ends at RNAi-mediated slice sites that allow specific adapter ligation and detection by RLM-RACE (B-C) or by NIL-TDS (B-D)) resolution and indicated turnaround times for each method are based on our experience and published data (Wang and Fang 2015). Only NIL-TDS generates quantitative outputs. Created with BioRender.com

In this study, we utilized the latest sequencing features to develop an innovative workflow for high resolution slice site detection in eukaryotes. We combined the ONT Flongle native ligation-based barcoding platform with RNA adapter ligation-mediated PCR approaches to establish <u>N</u>ative <u>I</u>ndex <u>L</u>igation-based <u>T</u>argeted <u>D</u>egradome <u>S</u>equencing (NIL-TDS) for the direct quantification of slicing activities of sRNA candidates (Fig. 1B-D). NIL-TDS offers higher sequencing depth as compared to RLM-RACE and PARE, which is most critical to detect slicing events in lowly expressed mRNAs. In addition, NIL-TDS is scalable, more specific, cost-effective, and offers a short turnaround time, while also being independent of tedious and bias-prone cloning. The high-read coverage and long-read capability of ONT enable precise detection of both high and low abundance slicing events. As a proof of concept, NIL-TDS was validated by analysing miRNA-induced slicing in plant seedlings and human cell lines. It revealed a previously unknown miRNA-mRNA cleavage-based interaction from a lowly expressed, highly relevant transcripts. The generated unprecedentedly detailed degradome profiles, quantified changes in sRNA slicing activity that improved our understanding of miRNA-mediated gene regulation.

## RESULTS

### NIL-TDS procedure outline

As a first step, an RNA adapter was ligated to the 5’-monophosphate ends of a sliced RNA, ensuring selective amplification of sRNA-effected cleavage products. This ligated RNA was converted to cDNA by reverse transcription, followed by PCR amplification using adapter-specific and target mRNA-specific primers to enrich for sliced mRNA amplicons. These amplicons were subjected to ONT Flongle sequencing using the native ligation-based barcoding platform, enabling multiplexing and parallel analysis on the same chip of samples or replicates, yielding more than 0.5 million reads per run. Sequencing reads were aligned to the reference transcriptome, and a bioinformatics pipeline was used to identify and quantify sRNA slicing sites. We developed an R script (see Supplementary Methods) for rapid data analysis. The script trims and aligns the base called and demultiplexed reads to the respective RNA to create high-resolution t-plots.

### Quantitative validation of the miR-196-*HOXB8* slice site

Human microRNAs (mir) are involved in almost every cellular process and their dysregulations have been shown to contribute to different human diseases including neurodegenerative diseases and cancer. Mir-196 is a conserved miRNA-family encoded within the HOX gene clusters and known to fine tune the expression of its target gene *HOMEOBOX B8* (HOX8B) and its aberrant expression is associated with tumerogenesis and development of lung metastases (Chen et al. 2011). To benchmark NIL-TDS sensitivity, we reanalysed the interaction of miR-196 with the *HOXB8* gene as the first discovered cleavage-mediated RNAi guide-target pair in mammals (Yekta et al. 2004). In their seminal study, Yekta et al. (2004) showed *HOXB8* cleavage activity with 8 RLM-RACE reads from total mouse RNA. Using NIL-TDS, we confirmed the expected mRNA cleavage at position 4918 as dominant signal (Fig. 2A), with ^211^/_3.368_ total reads in human lung carcinoma A549 cells which robustly expressed *HOXB8* (Jiang et al. 2024). Interestingly, additional and previously undescribed slicing signals were detected in both repetitions at positions 5,097, 5,123 and 5,144.

**Figure 2:**
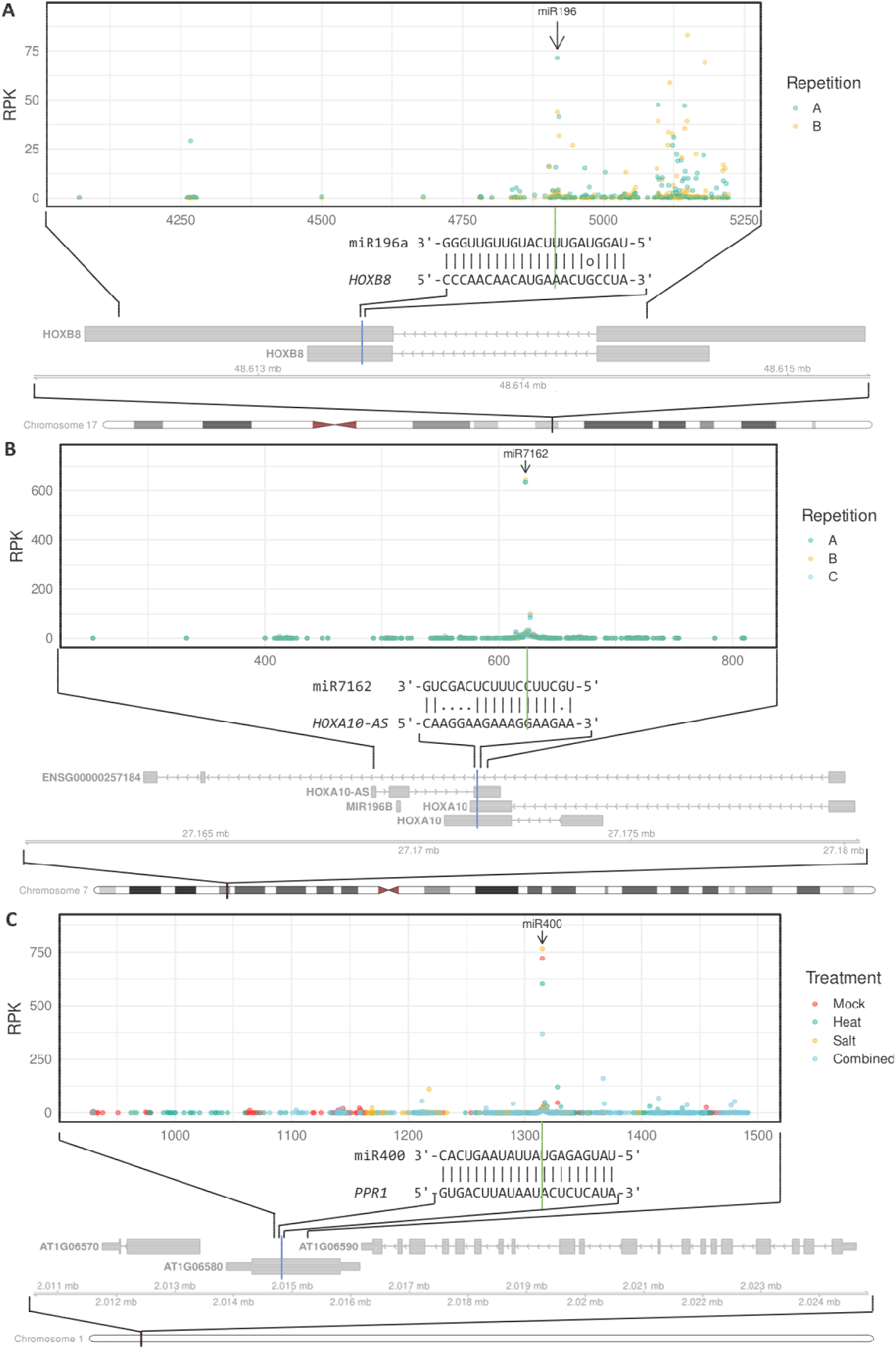
Quantitative NIL-TDS output in human and plant cells. NIL-TDS-based detection of HOXB8 slicing by miR-196 in human cells (A), HOXA10-AS slicing by miR-7162 in human cells (B), and PPR1 slicing by miR400 in Arabidopsis thaliana seedlings (C). Plotted positions are indicated on the genomic tracks, and blue lines mark respective miRNA target sites. Each point shows the number of reads per 1000 total reads, that aligned with their 5’- end to the respective mRNA base position indicating mRNA fragment ends. Arrows indicate the respective slice sites in the plots. Legends show colour-coded stress treatments or repetitions. The alignment between miRNA and target RNA is shown with a green line indicating slice-sites.

### Identification of a formerly unknown miR-7162-*HOXA10-AS* slice site with relevance in carcinogenesis

To test NIL-TDS capabilities to elucidate rare and potentially unknown slicing events, we investigated the complex and convoluted regulation of the HOX-gene cluster and its important regulator miR-196. One interesting precursor transcript of miR-196 is *HOXA10-AS* (antisense). HOXA10-AS is emerging as an key player in cancer biology, frequently overexpressed in several tumor types including glioma, breast, gastric and colorectal cancer. Its role in promoting cancer progression suggests its potential as a diagnostic and prognostic biomarker, making it a promosing target for therapeutic interventions (Hu et al. 2025). We were particularly interested in a potential miR-196 target site in this transcript, which would constitute a negative feedback loop to regulate miR-196 expression. Additionally, we wanted to investigate a potential regulatory mechanism that could decouple *HOX* gene expression from the suppressing influence of miR-196 which is co-transcribed from the same gene cluster (Hu et al. 2025). Therefore, we generated three replicates of NIL-TDS with a total of 64,350 reads from the *HOXA10-AS* transcript (Fig. 2B) in A549 cells. Surprisingly, we identified one highly consistent degradation signal at position 623, with 647, 638 and 636 reads per 1000 reads (RPK). This position corresponded to the predicted slice site of miR-7162, a yet undescribed interaction, indicating suitability of NIL-TDS for discovering previously unknown miRNA activities (Fig. 2B). In the course of our studies, we anticipated that NIL-TDS might introduce PCR duplication events in sRNA targets resulting in an overestimation of slicing. To quantify the extent of PCR duplication, HOXA10-AS was reanalysed with the incorporation of unique molecular identifiers (UMIs). In a first step, random RNA heptameres were ligated to RNAs, to serve as UMIs, followed by 5’ phosphorylation and RNA adapter ligation. We observed that this UMI approach incorporated unspecific 5’ phosphorylation, leading to the amplification of all RNAs and reducing the fraction of reads mapping to HOXA10-AS to 13 % (2472 / 18,460). However, the obtained 2,472 reads represented 371 unique barcodes with a saturating sequencing depth and a dominant and statistically approvable slicing signal at position 623 (Fig. S1).

### Detection of climate stress-induced minute changes in miR400-mediated slicing of PPR1

To validate the applicability and sensitivity of NIL-TDS across eukaryotes, we analysed the slicing site of *Arabidopsis thaliana* miR400 (*Ath*-miR400) and its target *PENTATRICOPEPTIDE REPEAT 1* (*PPR1*; *AT1G06580*). *Ath*-miR400 is downregulated (Barciszewska-Pacak et al. 2015) while *PPR1* is upregulated during various stresses, such as salt and heat stress (Kilian et al. 2007). However, the hypothesised reduction in slicing activity has never been experimentally validated. Therefore, we treated *Ath* seedlings with 250 mM NaCl (salt stress), without NaCl (mock), exposed them to 37°C (heat stress), or subjected them to a combination of salt and heat stress (combined stress). In total, we generated 214,087 reads, of which 154,786 fulfilled the quality criteria to carry the ligated slicing-adapter and to align end-to-end with at least 100 nt to the target transcript (Tab. S1). NIL-TDS provided an unprecedented resolution of the *Ath*-miR400 target site degradation profile on *PPR1* (Fig. 2C). Confirming NIL-TDS quantitative sensitivity. We observed a slight reduction of slicing activity upon heat stress, which was significantly attenuated upon combined stress treatments (Fig. 2C, Fig. S2).

### NIL-TDS is precise and reproducible, but amplifies rare exosomal degradation events in the absence of targeted slicing

To further evaluate the precision and reproducibility of NIL-TDS, we conducted two control experiments. First, we measured NIL-TDS accuracy by amplifying a 590 bp long amplicon (from outer amplicon of 691 bp) of *PPR1* cDNA from each *Ath* sample. Over 80% of all reads mapped to the respective inner primer binding site or 1 nt downstream, while 4% mapped to the outer amplicon (Fig. 3). This demonstrated the precision (Fig. 3A) and the reproducibility between replicates (Fig. 3B) across PCR, sequencing and bioinformatics.

**Figure 3:**
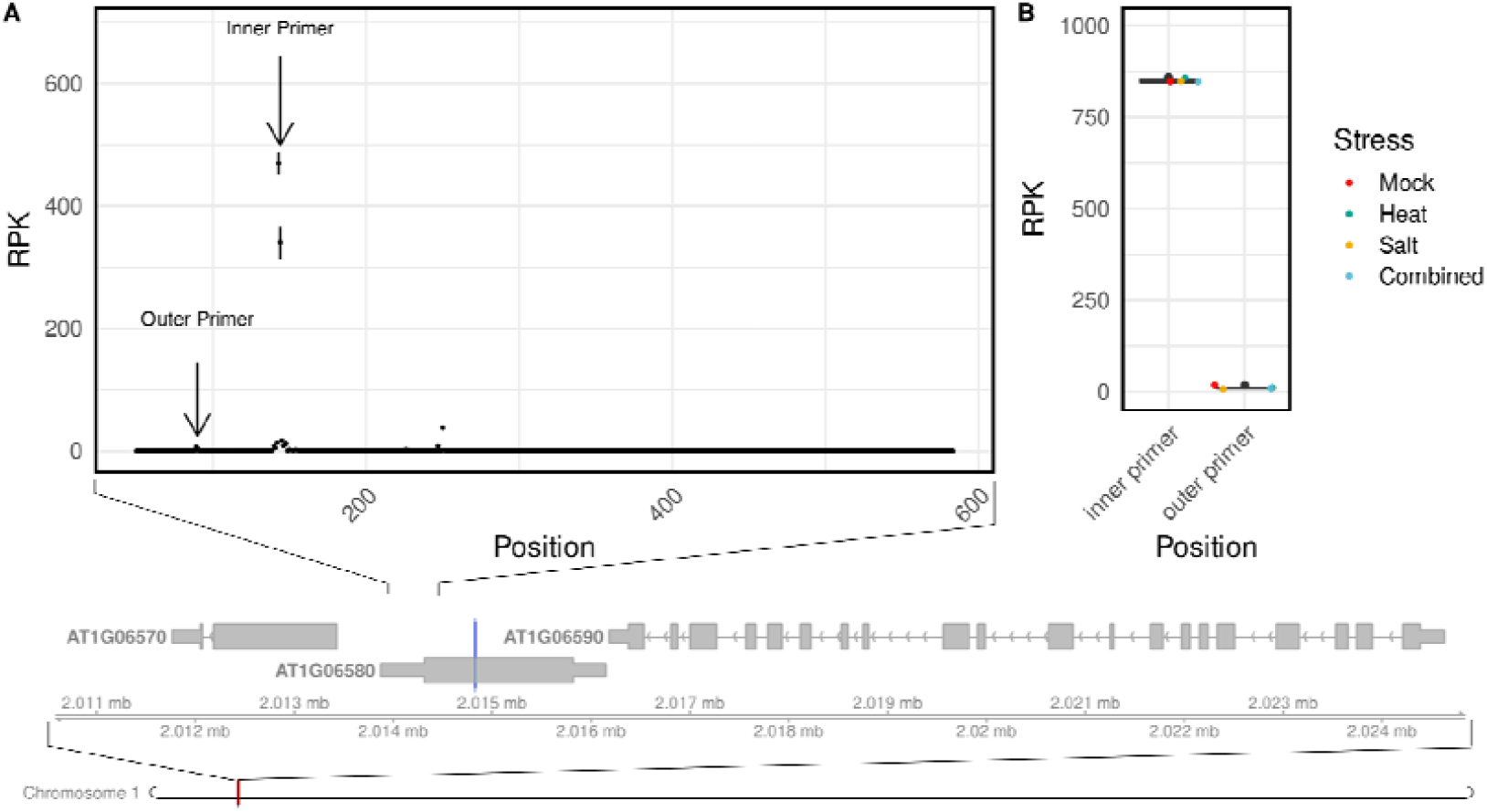
Precision of NIL-TDS on a cDNA amplicon. A nested PCR was conducted using *A. thaliana* Col-0 cDNA for the amplification of *PPR1* amplicon from four different stress treatments. Amplicons were sequenced on a nanopore Flongle cell. Start sites of alignments (A). Fraction of reads within the primer start site +/-2 bp (B). Each point shows the number of reads per 1000 total reads, that aligned with their 5’-end to the respective mRNA base position indicating mRNA fragment ends. Bars show the standard deviation between the treatments. Plotted positions are indicated on the genomic tracks, and a blue line marks the miRNA target site.

Secondly, we applied NIL-TDS to a region of *PPR1* with no known slicing sites (Fig. 4). Some positions showed a slicing signal independent of the different stress treatments, which we interpreted as unspecific exonucleolytic degradation. In fact, these sites were clearly distinguishable from true slicing events by analysing replicates of nested PCR reactions. Accurate and specific slicing events typically exhibited consistent read patterns corresponding to the known miRNA target sites, while unspecific degradation showed a random distribution of read points, which were not detected in replicates.

**Figure 4:**
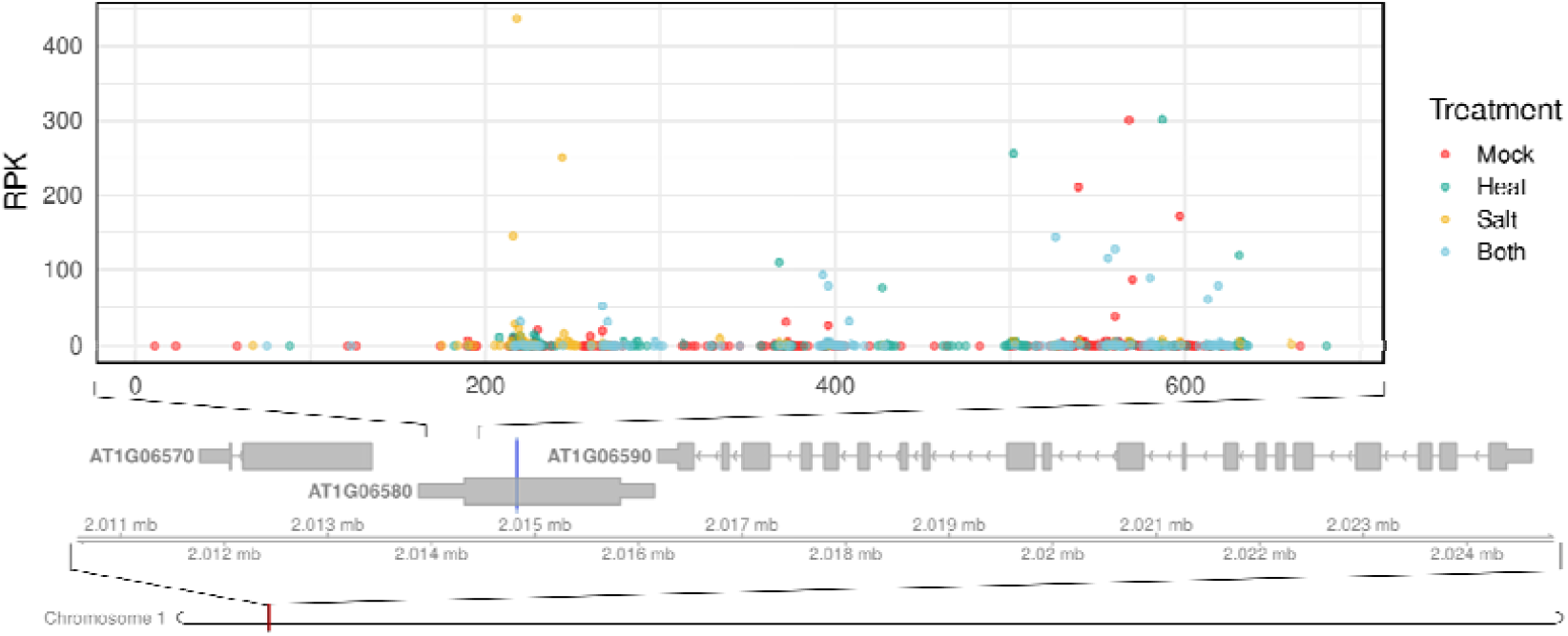
Comparative NIL-TDS readout of a non-target site of *PPR1* under abiotic stress. NIL-TDS readout of an untargeted site of *PPR1*. Each point shows the number of reads per 1000 total reads, that aligned with their 5’-end to the respective mRNA base position indicating mRNA fragment ends. Stress treatments are indicated by colour (mock (red), heat (green), salt (yellow), and combined (blue)). Plotted positions are indicated on the genomic tracks, and blue lines mark respective miRNA target sites. Positions indicate the first base after the slice-site, which is also the first base after the 5’ RLM-adapter.

Consistent with this, in all experiments, the mean predicted sequencing error rate was below 0.001 (≥ Q30) as indicated by the base calling algorithm, and the median relative alignment edit distance was below 3%.

## DISCUSSION

### Revisiting miR-196 slicing activity reveals high sensitivity of NIL-TDS

Our study indicated NIL-TDS as a highly sensitive and accurate tool for mapping sRNA slicing sites and quantifying their activity. Integrating nanopore sequencing, NIL-TDS generated up to 100,000 times more reads than current standard methods (RLM-RACE, PARE), facilitating the detection and statistical validation of abundant to rare slicing events. Compared to RLM-RACE, it offers higher scalability and precision while providing a streamlined, ready-to-use workflow with broad applicability in basic and applied life sciences. As a validation for the methodology, we quantified the degree of slicing in the first ever discovered RNA-guided slicing event in mammalian systems. The detection of miR-196-mediated slicing of *HOXB8* had been a challenging task in past studies. Bracken et al. (2011) could not detect slicing of *HOXB8*, despite using mouse embryonic tissue, as used by Yekta et al. (2004) in their seminal study, via PARE. While Shin et al. (2010) were able to show *HOXB8* slicing with a PARE approach in HeLa cells, their approach produced only two reads of *HOXB8* and only one read was located at the expected slice site. This highlights the difficulty of detecting slicing events for genes with low expression, like *HOXB8* in HeLa cells (Sjöstedt et al. 2020), with current methods.

We selected A549 cell lines, known for their more robust expression of *HOXB8*. Due to the inverse correlation of gene expression and slicing activity, the selection of a cell line with robust *HOXB8* expression for our experiments carried the risk of undetectable slicing activities. In contrast, the selection of HeLa cell lines, by Shin et al. (2010), with a diminished *HOXB8* expression generated barely sufficient reads. Therefore, we expected a lower fraction of sliced *HOXB8* transcripts in A549 as compared to HeLa cells. The low percentage of only 6% (in contrast to 50% in HeLa cells detected by Shin et al. (2010) of all reads at the slice site of miR-196 confirmed this expectation. Due to the higher sensitivity of NIL-TDS, we were still able to robustly detect the very rare slicing event.

### Significance of a new slice site in a miR-196-precursor

*HOX* genes have pro-oncogenic functions in many types of cancers (Bhatlekar et al. 2014; Morgan et al. 2022). Surprisingly, miR-196, which can downregulate a fourth of all *HOX* genes, also has pro-oncogenic effects and is regarded as a potential cancer biomarker (Khalilian et al. 2024). This motivated us to analyse the biogenesis of miR-196b, which originates from the *HOXA10-AS* transcript, and we found that it involves miR-7162. The expression of miR-7162 is undetectable in all cell lines in miRmine (a database of human miRNA expression profiles) (Panwar et al. 2017), and therefore, miR-7162 activity remained elusive. The previously unknown slicing activity of miR-7162-5p detected in our study might explain reduced miR-196 expression in the A549 cell and other cell lines or tissues. Importantly, our findings represent a promising starting point for future functional analyses of miR-196 and miR-7162. The sensitivity and accuracy of NIL-TDS can be particularly valuable in such studies. Recent studies suggested that miR-7162-3p, delivered via exosomes from human umbilical cord mesenchymal stem cells, plays an important role in repairing endometrial stromal cell injuries by targeting and restricting *Apolipoprotein L6* (*APOL6*), a protein associated with tumour aggressiveness, thereby reducing cell death and inflammation (Shi et al. 2021). Furthermore, Chakraborty and Nath (2022) reported miR-7162-5p to potentially target *COL5A1*, a gene encoding the alpha-1 chain of type V collagen, which is associated with malignancy of certain cancers, including lung cancer, promoting tumour progression and metastasis. The latter study suggested that the absence of miR-7162-5p may contribute to oncogenic processes by regulating *COL5A1* expression, thereby potentially enhancing cancer cell proliferation and metastasis. NIL-TDS could validate this role of miR-7162-5p in healthy tissues. Therefore, further analyses of miR-7162 mimics (synthetic miRNAs behaving similarly to endogenous miRNAs) and NIL-TDS based analyses may elucidate the role of miR-196 in cancer development and potentially open new avenues for miRNA-based drugs and therapeutics.

### Stress-induced changes of *Ath-*miR400 slicing activity explain *PPR1* induction

Different studies have analysed the expression of *Ath*-miR400 and its target gene *PPR1* upon abiotic stress in *A*. thaliana (Barciszewska-Pacak et al. 2015; Kilian et al. 2007). Overexpression of *Ath*-miR400 results in an increased susceptibility of *A. thaliana* plants to bacterial and fungal pathogens (Park et al. 2014). Commonly, these studies rely on multi-omics approaches, correlating sRNA and mRNA expression patterns and determining target cleavage via RLM-RACE. To validate the causative relation between sRNA and mRNA expression, sRNA overexpression lines along with mRNA knockout mutants are often analysed by RLM-RACE. To demonstrate that this tedious endeavour can be circumvented with NIL-TDS on stressed plants, we analysed the slice site of *Ath*-miR400 under various stress conditions. Our analysis generated, for the first time, quantitative data that confirmed previous qualitative results. Importantly, this quantitative NIL-TDS read-out of *Ath*-miR400 activity might elucidate the significance of target transcript abundances and turnover in plant adaptative signalling to climate stress. It suggests a regularity model in which RNAi-based slicing finetunes transcript levels at high spatio-temporal tissue resolution and compensate for the regulatory less dynamic mRNA biogenesis machinery.

### Potential of NIL-TDS for quantitative slicing measurements

Several poorly understood parameters can influence the effectiveness of sRNAs (Diener et al. 2023), while off-target effects pose significant challenges in mode-of-action studies of therapeutics and plant protectants. At the same time, computational target prediction algorithms can identify a plethora of potential off-targets (Kern et al.2019), and bioinformatic dsRNA design tools can minimize but not completely avoid off-target activities (Ahmed et al. 2020). Off-target alleviation is often attempted by using sRNA pools, either through the usage of long dsRNA, which can produce an array of siRNAs, or by the design of up to 30 different sRNAs as an effort to “dilute” off-target effects (Ahmed et al. 2020; Neumeier and Meister 2021). The so called “too many targets for miRNA effect” (TMTME) highlights the prevalence of these issues (Zhang et al. 2021). NIL-TDS can address underlying challenges by detecting even rare off-target events. The generated data sets can be used to further improve target prediction algorithms. In terms of application, NIL-TDS can assist in the development of RNA-based therapeutics and plant protection products by assessing the efficiency of multiple formulations and different degrees of sRNA-target complementarity.

### Excessive PCR cycles overamplify exonuclease degradation

The application of NIL-TDS to an untargeted region of *PPR1* revealed seemingly random peaks in the degradomes of different samples, despite the demonstrated precision of the technique. Such incidences are, to our opinion, the result of the amplification of undirected exonuclease degradation products, which also carry monophosphate-ends. In our efforts to distinguish between true and false signals we successfully developed several strategies. For the de-novo discovery of miR-7162-mediated slicing, we conducted multiple independent replicates, which showed a consistent slicing signal, confirming targeted slicing activity. Additionally, in the case of miR400 and miR-7162, target genes showed a dominant slice site, indicating targeted exonucleolytic cleavage. In the cases of miR-196- and miR400-mediated slicing we could found our conclusions, additionally to the independent replications, on prior knowledge. The most reliable strategy to distinguish true slicing events from random exonuclease products is to perform biological replicates, ensuring that reproducible cleavage sites reflect directed endonuclease activity. Repeating nested PCR reactions across independent samples helps confirm the consistency of true slice sites. While the directed endonuclease slice sites are reproducible, undirected exonuclease products are not. For instance, German et al. (2008) alleviated a similar challenge by knocking-out the responsible nuclease *XRN1* in *A. thaliana* (Benesova et al. 2021).

### Flongle provides optimal throughput and read lengths for targeted degradome sequencing

Dynamic development of sequencing technologies provided platforms varying in cost efficiency and throughput. For NIL-TDS, we considered different approaches on different sequencing platforms. We aimed at 50,000 reads per sample to obtain sufficient read-depth saturation for individual targets. Compared to the ONT Flongle, other platforms have critical drawbacks. While Illumina sequencing by synthesis technology reads are cost competitive on the highest throughput devices (Tab. S2), it faces limitations with long templates. In addition, obtaining short read depths of 50K reads per sample at competitive costs using high throughput NextSeq2000 flow cells would involve the almost infeasible multiplexing of 36K samples (Tab. S2). Additionally, NIL-TDS on Flongle is twice as fast as the iSeq100, and the low cost of the sequencer device makes in-house sequencing affordable and convenient. Together with a simpler and faster library preparation the NIL-TDS workflow is currently the most time and cost-effective solution for targeted slice site analyses in specific transcripts (Fig. 1).

To conclude, NIL-TDS represents notable progress in quantifying sRNA slicing activity and offers an unprecedented increase in read depth compared to traditional methods like RLM-RACE. This increased sensitivity enables the identification of rare slicing events, as demonstrated in the identification of miR-7162-*HOXA10-AS* slice-site, and confirming miR-196-*HOXB8* interactions in human lung carcinoma cells. The accuracy and sensitivity of the method in different eukaryotes was also confirmed by validating minute changes in stress-induced miRNA-mRNA interactions in *Arabidopsis thaliana* quantifying *Ath*-mir400 slicing of PPR1. By utilising the ONT flongle flow cell, NIL-TDS offers a fast, accurate and cost-effective solution for in-house transcript-specific degradation analyses, making it a scalable and competitive alternative to conventional methods.

## MATERIAL AND METHODS

### Plant material and stress treatment

*Arabidopsis thaliana* Columbia-0 (Col-0) seeds were grown on vertical square Petri dishes on ½ MS (Murashige and Skoog) media for 14 days in a 22°C day/18°C night cycle (8 h of light) preceded by 2 days stratification at 4°C. Complete seedlings were harvested and placed in a flask with ½ MS liquid media. Heat-treated plants were subject to 37°C temperature for two hours. For salt stress treatment, NaCl concentration was adjusted to 250 mM. Seedlings exposed to a combination of heat and salt stress contained 250 mM NaCl and were subject to 2 h heat at 37°C. Mock seedlings were incubated for 2 h in ½ MS liquid culture and put back in the growth chamber.

### Human cell line

A549 (ECACC86012804) cells were grown in DMEM containing Glucose, GlutaMAX and pyruvate supplemented with 100 U/ml penicillin, 100 µg/ml streptomycin and 10% (v/v) fetal bovine serum (all from Thermo Fisher Scientific, USA). The cells were cultured at 37°C and 5% CO_2_ in a humidified atmosphere.

### RNA extraction from plant material

Whole Arabidopsis seedlings were harvested from ½ MS liquid medium, briefly dried on a soft tissue, shock-frozen in liquid nitrogen, and processed to powder using a tissue lyser (Qiagen Retsch TissueLyser II). Total RNA was extracted using Direct-zol RNA Miniprep (Zymo Research, R2051) according to the manufacturer’s instructions with on-column DNase I treatment. RNA concentration was determined using NanoDrop ND-1000 spectrophotometer (Thermo Fisher Scientific, USA), and purity was determined by measuring A260/280 and A260/230 ratios. RNAs were stored at -80°C until further use.

### RNA extraction from human cells

For cell lysis, 5 ml of Trizol was added to the human cell line (A549) pellet (50×10^6^ cells), and 1 ml was taken for forward RNA extraction. The lysed cells were incubated at room temperature for 5 minutes and then centrifuged at 16,000x g for 1 minute to remove cell debris. The supernatant was transferred to a new tube. 200 µl of chloroform was added, and the cells were vortexed vigorously for 15 seconds, incubated at room temperature for 3 minutes, then centrifuged at 12,000x g for 15 minutes at 4°C. The upper aqueous phase containing RNA was transferred to a new tube. 2x volume of cold 100% ethanol was added, and the sample was incubated overnight at -20°C and then centrifuged at 12,000x g for 10 minutes at 4°C. The supernatant was removed, and RNA pellet was washed with 1 ml of 75% ethanol. The sample was mixed by vortexing and centrifuged at 7500x g for 5 min at 4°C. The washing step was repeated, and all ethanol residues were removed. The pellet was air-dried, and the RNA was dissolved in 50 µl DEPC water.

### RNA adapter ligation

Two µg of RNA isolated from both *Ath* plants and A549 human cells were used as a template for ligation with 2 µl 5′RLM-RACE adapter (5’-GCUGAUGGCGAUGAAUGAACACUGCGUUUGCUGGCUUUGAUGAAA-3’) [0.3µg/µl] (Eurofins Genomics, Germany). The ligation was performed using T4 RNA Ligase [10 U/µL] and its accompanying reagents (Thermo Fisher Scientific, USA). The reaction mixture contained 1 µl of 10X Reaction buffer, 1 µl BSA [1 mg/ml], 1 µl of T4 RNA ligase, and DEPC water, adjusted to a final volume of 10 µl. The reaction mixture was incubated for 60 min at 37°C in a thermocycler, following Wang et al. (2015). All components were added individually and not in a master mix, with the RNA and the RLM adapter being added last.

### Reverse transcription

The total ligated RNA (10 µl) was used directly to generate the first cDNA strand using the RevertAid Reverse Transcriptase (Thermo Fisher Scientific, USA). A master mix was prepared for cDNA generation containing 1 µl of random hexamers [100 pmol/µl], 4 µl of 5X Reaction buffer, 0.5 µl of Ribolock RNase inhibitor, 2 µl of dNTP mix [10 mM], and 1 µl of RevertAid reverse transcriptase (Thermo Fisher Scientific, USA). The volume was adjusted to 20 µl with water. The reaction was run at 25°C for 10 min, 42°C for 60 min, and 70°C for 10 min.

### UMI-RLM adapter ligation and reverse transcription

2 µg RNA from A549 human cells were first ligated to a Unique Molecular Identifiers (UMI) adapter (Random RNA Heptamer: 5′-NNNNNNN-3′) [0.3 µg/µl] (Integrated DNA Technologies, USA) in a 10 µl reaction using T4 RNA ligase following the protocol previously described for the RLM adapter ligation. The total UMI ligated RNA product was phosphorylated using 1 µl [10 U/µl] T4 Polynucleotide Kinase Ligation (T4 PNK, New England Biolabs, M0201S), 4 µl PKN Buffer and 5 µl dATPs [10 mM] in a total volume of 10 µl. The mixture was incubated at 37°C for 30 min, followed by an inactivation step at 65°C for 20 min. The phosphorylated UMI-ligated RNA was purified using the RNA clean and concentration kit (Zymo Research, R1017) and eluted in 10 µl nuclease-free water. The entire eluate was then used for RLM-adapter ligation using T4 RNA ligase as previously described in a 15 µl reaction. The resulting UMI-RNA-RLM adapter product was subsequently used to generate the first cDNA strand. Reverse transcription was carried out in a total volume of 25 µl, replacing random hexamers with Oligo dT (16x, 18x, 20x) primer [100 pmol/µl] and in a total volume of 25µl.

### Nested PCR

Two rounds of nested hot-start touch-down PCR were performed using outer (first) and inner (second) 5′-RLM-RACE universal primers in combination consecutively with gene outer specific (e.g., PPR1_Target_Outer) and gene inner specific (e.g. PPR1_Target_Inner) primers. GoTaq® G2 Hot Start Master Mix (Promega, M7432) was used for the nested PCR. A master-mix was prepared for both outer and inner PCRs, containing 25 µl of 2X GoTaq® G2 Hot Start Master Mix, 1.5 µl of RLM-Universal Outer/Inner Primer [10 pmol/μl], 1.5 µl of RLM-Outer/Inner-Specific Primer [10 pmol/μl], and 2.5 µl of cDNA or prior PCR product, with the total volume adjusted to 50 µl with water. As a control, PPR1_Non-Target_Outer and PPR1_Non-Target_Inner primers were used for amplification of a non-target site of *PPR1*, replacing PPR1_Target_Outer and PPR1_Target_Inner primers in the reaction. The thermocycler was preheated at 95° C and samples were denatured at 95°C for 5 minutes followed by 18 touch-down cycles of denaturation at 95°C for 30 sec, annealing at 68°C (incrementally decreased by 0.5°C per cycle) for 30 sec, and extension at 72°C for 30 sec. After the first 18 cycles, 20 additional cycles of 30 sec denaturation at 95°C for 30 seconds is performed, followed by annealing at 62°C for 30 seconds and extension at 72°C for 30 seconds. The final extension step is 72°C for 5 minutes, and the reaction is then held at 4°C.

For the human cell line (A549), 3.5 µl cDNA or PCR product from the prior outer PCR reaction was used as a template for the nested PCR. For the human cell line (A549) initial annealing temperature for the touchdown PCR was optimised for 66°C. Inner and outer PCR products were evaluated in a 1% agarose gel and bands from 100 to 1000 bp were excised. Products were cleaned with the Wizard^®^ SV Gel and PCR Clean-Up System (Promega). DNA concentration was measured using Qubit™ dsDNA HS Assay Kit (ThermoFisher Scientific, USA; Invitrogen, USA) and purified PCR products were stored at -20°C until further use. All used oligonucleotides are listed in Table 1.

**Table 1:**
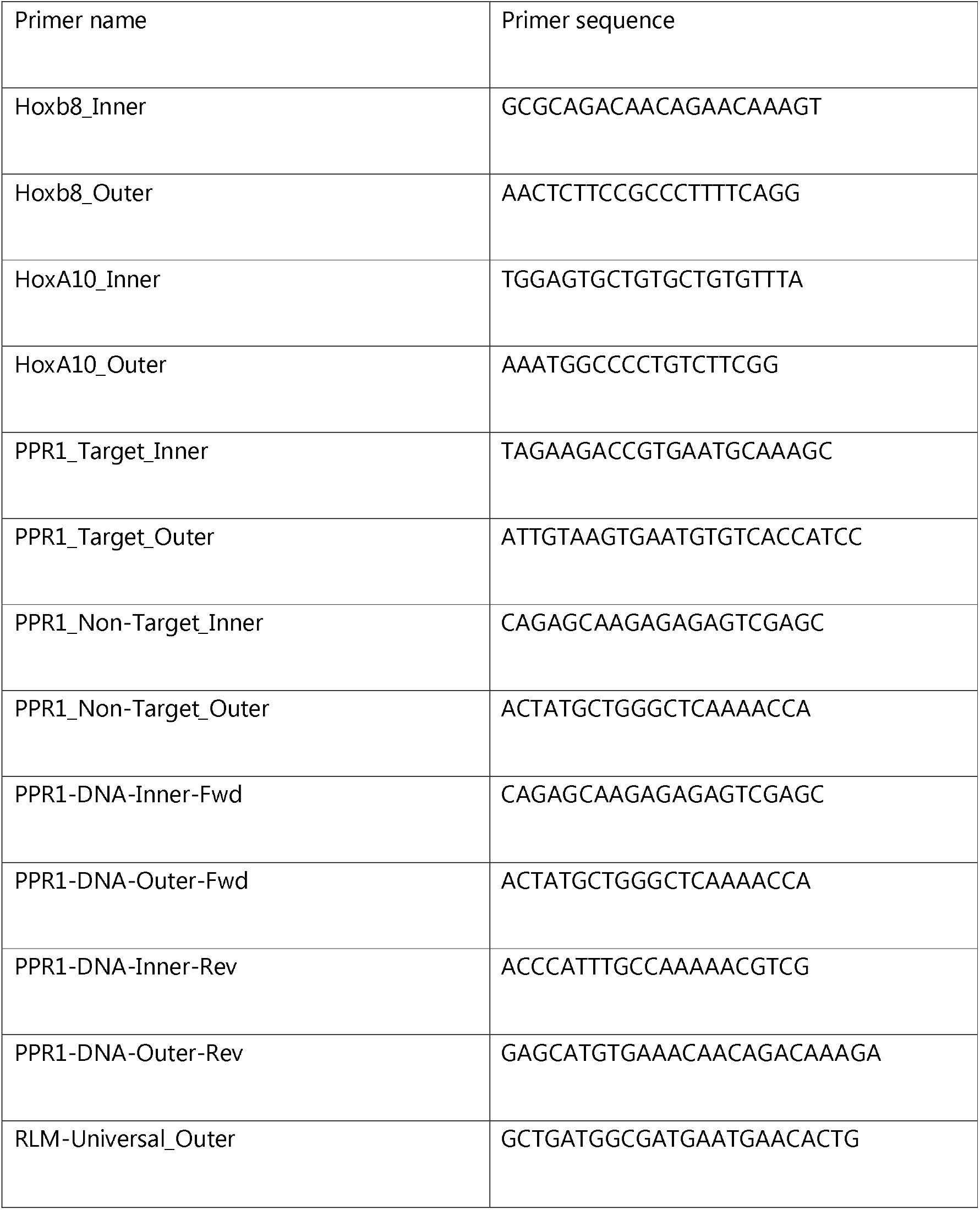
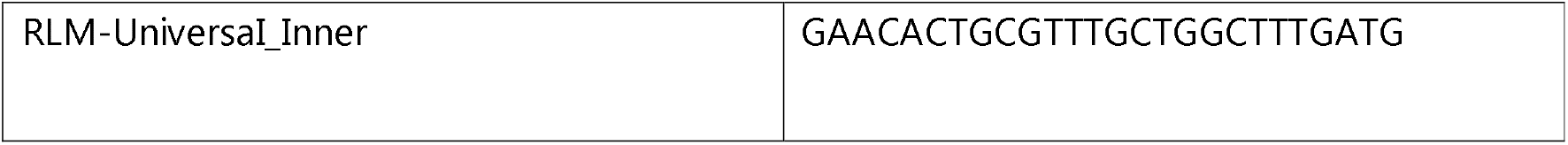
Primer list.

### PCR amplification for cDNA control

Two µg of the RNA extracted from *Ath* plants (mock and stress treated) were used for reverse transcription (as previously described) without an adapter ligation step. As a control, two rounds of hot-start touch-down PCR were performed using PPR1_DNA-outer-Fwd and Rev primers (first round) and PPR1_DNA-Inner-Fwd and Rev primers (second round) for the amplification of a *PPR1* amplicon. The GoTaq® G2 Hot Start Master Mix (Promega, M7432) was used for the nested PCR. A master mix was prepared for both outer and inner PCRs. The assembled reaction and the thermocycler program are as described in the nested-PCR section using the mentioned primers (Tab. 1).

### Sequencing

Libraries were prepared for 12 - 16 PCR products according to the Ligation sequencing amplicons - Native Barcoding Kit 24 V14 protocol (SQK-NBD114.24) (https://community.nanoporetech.com/docs/prepare/library_prep_protocols/ligation-sequencing-amplicons-native-barcoding-v14-sqk-nbd114-24/) of Oxford Nanopore Technologies (ONT). Libraries were run on an ONT MinION Mk1B device with a flongle adapter on a Flongle Flow Cell (R10.4.1). Libraries were loaded according to Russel (2023). Sequencing data was recorded for 24h with a length cut-off of 20bp as pod5 data.

### Bioinformatics

#### Sequencing, trimming & alignment

Sequencing raw data was basecalled via the ONT dorado (v0.5.2+7969fab) with a minimal q-score of 12, the kit option SQK-NBD114-24 and the Super Accuracy (“sup”) basecalling algorithm. Reads were not trimmed, and a fastq-file was emitted which was subsequently demultiplexed. Quality control was conducted via FastQC v0.12.1 (Andrews 2010). To remove adapters and orient the reads, the RLM-adapter sequence from inner primer binding site to adapter-3’-end (TTTCATCAAAGCCAGCAAACGCA) was used to trim reads with cutadapt v4.2 (Martin 2011) with Python 3.11.6 (Van Rossum and Drake 2018). Correct orientation of reads was ensured by options “--rc” and “--trimmed-only”. Trimmed reads were aligned with bowtie2 v2.5.0 (Langmead and Salzberg 2012) in end-to-end mode and score-min set to “L,0,-0.6”. Unique molecular identifiers (UMIs) were extracted before alignment and alignments were deduplicated with umi_tools v1.1.6 (Smith et al. 2017).

#### Analysis & plotting

To analyse the alignments, the bowtie pseudo sam-files were read into R (Team 2014) v4.3.1 via read.delim. Alignments were filtered by flag 16 and afterwards, the alignment start position indicates the slice site. Results were plotted with ggplot2 (Wickham 2024) v3.4.3, ggpubr (Kassambara 2024) v0.6.0 using the palettes of wesanderson (Ram et al. 2024) v0.3.6. An example script is provided as R-Script S1. Genomic tracks were created with Gviz (Hahne and Ivanek 2016) v1.46.1. Track information was either downloaded from UCSC genome browser (Lee et al. 2019) for the human chromosomes (hg38, GCF_000001405.40) via the UcscTrack function or from NCBI RefSeq (O’Leary et al. 2015) for the *A. thaliana* chromosome (GCF_000001735.4) via the makeTxDbFromGFF function from GenomicFeatures (Lawrence et al. 2013) v.1.54.4.

## Supporting information

NIL-TDS supplementary

## ABBREVIATIONS

NIL-TDS: Native index ligation-based targete degradome sequencing
RLM-RACE: RNA ligase-mediated rapid amplification of cDNA ends
PARE: parallel analysis of RNA ends
HOX: *HOMEOBOX*
ONT: Oxford Nanopore Technology
A549: Human lung carcinoma cells

## DECLARATIONS

### Availability of data and materials

The data underlying this article are available in the EMBL-EBI European Nucleotide Archive (ENA) at https://www.ebi.ac.uk/ena/, and can be accessed with PRJEB85122. Direct access to sequencing data is provided via: https://ftp.sra.ebi.ac.uk/vol1/run/ERR141/ and the accessions in Table S1.

### Authors′ contributions

Bernhard T. Werner: Conceptualization, Data curation, Formal analysis, Investigation, Methodology, Project administration, Software, Resources, Supervision, Validation, Visualization, Writing – original draft, Writing – review & editing. Sabrine Nasfi: Conceptualization, Funding acquisition, Investigation, Methodology, Resources, Validation, Writing – original draft, Writing – review & editing. Jens Steinbrenner: Conceptualization, Methodology, Writing – review & editing. M. Lienhard Schmitz: Investigation, Resources, Writing – review & editing. Manar Makhoul: Investigation, Methodology, Resources, Writing – review & editing. Patrick Schäfer: Conceptualization, Funding acquisition, Project administration, Resources, Supervision, Writing – original draft, Writing – review & editing

## Acknowledgements

We want to thank Markus Schwinn for providing materials and Marek Bartkuhn for scientific discussions.

## SUPPLEMENTARY DATA

Supplementary Data will be available on RNA journal online upon publication.

## TABLE AND FIGURES LEGENDS

Figure S1. Effect of deduplication via UMIs. To assess the effect of PCR duplicates on NIL-TDS UMIs were incorporated in the workflow and reads were deduplicated with UMI_tools. Due to PNK1 phosphorylation during library prep, 2,147 reads of the 18,460 generated reads aligned to HOXA10-AS. After deduplication 241 reads remained, confirming prior results. Each point shows the number of reads per 1000 total reads, that aligned with their 5’-end to the respective mRNA base position indicating mRNA fragment ends. The arrow indicates the respective slice site of miR-7162. Colour indicates results before (yellow) and after (blue) deduplication.

Figure S2. NIL-TDS-based detection of slicing by miR400 in *PPR1* (Repetition). Plotted positions are indicated on the genomic tracks and a blue line marks the miRNA target site. The number of reads per 1000 total reads (RPK) of total reads corresponding to the respective position of the target RNA are shown. The respective slice sites are indicated by arrows and text. Stress treatments are indicated by colour. The alignment between miRNA and target RNA is shown and a blue line indicates the slice-site.

Table S1: Sequencing statistics related to the samples, their quality, read lengths, sequencing depth and the ENA accessions. The table gives information for each library. Total reads are reads for the respective library after demultiplexing. Mean Q is the mean of means of - log_10_(error rate). SD stands for standard deviation and L for read length. Filtered reads shows the number of reads with a proper RNA adapter.

Table S2: Cost and throughput of different sequencing platforms. The here presented costs are the list prices from the manufacturers homepages (January 2025).

RScript S1. A script to easily evaluate NIL-TDS data and to produce t-plots.

