## Supplementary material for "High Precision Quantification of small RNA Slicing Activity - Native Index Ligation-based Targeted Degradome Sequencing (NIL-TDS)": NIL-TDS supplementary

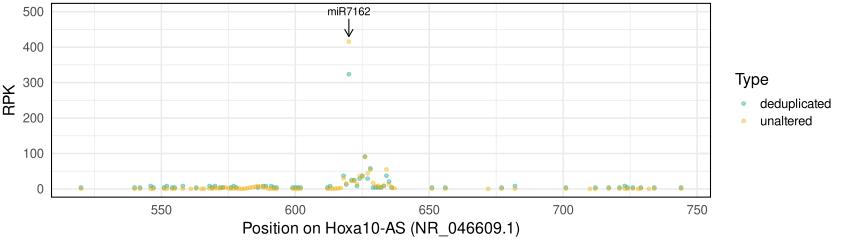

Figure S1. Effect of deduplication via UMIs. To assess the effect of PCR duplicates on NIL-TDS UMIs were incorporated in the workflow and reads were deduplicated with UMI_tools. Due to PNK1 phosphorylation during library prep, 2,147 reads of the 18,460 generated reads aligned to HOXA10-AS. After deduplication 241 reads remained, confirming prior results. Each point indicates the number of reads per 1000 total reads, that aligned with their 5’-end to the respective mRNA base position and therefore indicate mRNA fragment ends. The arrow indicates the respective slice site of miR-7162. Colour indicates results before (yellow) and after (blue) deduplication.

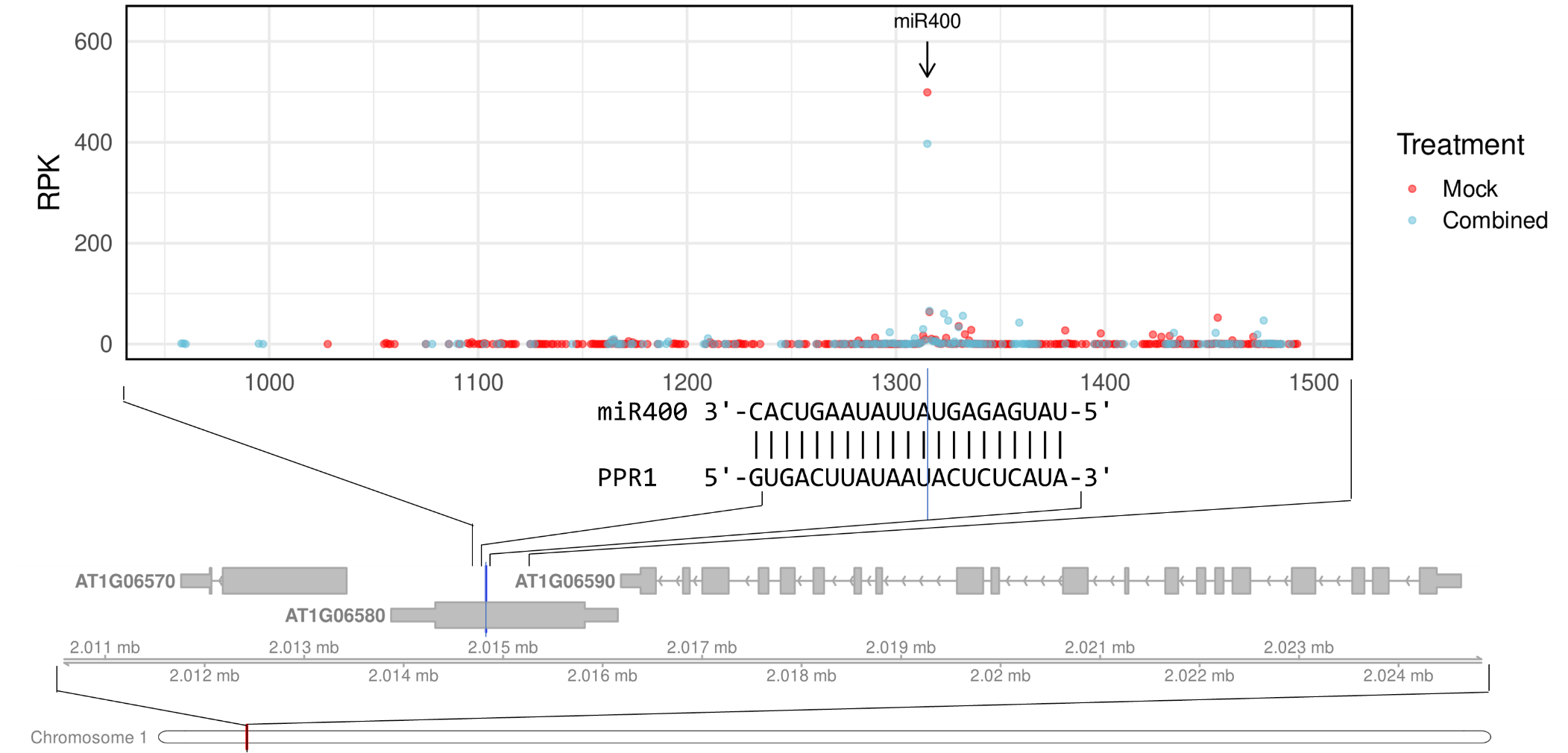

Figure S2. NIL-TDS-based detection of slicing by miR400 in *PPR1* (Repetition). Plotted positions are indicated on the genomic tracks and a blue line marks the miRNA target site. The number of reads per 1000 total reads (RPK) of total reads corresponding to the respective position of the target RNA are shown. The respective slice sites are indicated by arrows and text. Stress treatments are indicated by colour. The alignment between miRNA and target RNA is shown and a blue line indicates the slice-site*.*

Table S1: Sequencing statistics related to the samples, their quality, read lengths, sequencing depth and the ENA accessions. The table gives information for each library. Total reads are reads for the respective library after demultiplexing. Mean Q is the mean of means of -log_10_(error rate). SD stands for standard deviation and L for read length. Filtered reads shows the number of reads with a proper RNA adapter. Data can be downloaded via: https://ftp.sra.ebi.ac.uk/vol1/run/ERR141/

| Amplification | Target region | ENA Accession | Total reads | Mean Q | SD(Q) | Mean L | SD(L) | Filtered reads | Filtered reads [%] | Trimmed L | SD (Trimmed L) | Trimmed mean Q | SD (Trimmed Q) | Alignments ≥ 100nt | Alignments [%] | Target site ±3 [%] |
| --- | --- | --- | --- | --- | --- | --- | --- | --- | --- | --- | --- | --- | --- | --- | --- | --- |
| RLM-RACE (inner) | miR400 target site | ERR14185211 | 54,043 | 32.9 | 4.66 | 317.14 | 156.24 | 46,093 | 85.29% | 284.97 | 148.76 | 33.07 | 4.9 | 41,120 | 76.09% | 74.54% |
| RLM-RACE (inner) | miR400 target site | ERR14185215 | 60,171 | 32.94 | 4.66 | 282.8 | 134.53 | 51,147 | 85.00% | 248.64 | 127.02 | 32.96 | 4.92 | 46,050 | 76.53% | 65.65% |
| RLM-RACE (inner) | miR400 target site | ERR14185202 | 41,975 | 32.98 | 4.77 | 295.51 | 144.7 | 35,628 | 84.88% | 256.87 | 141.55 | 32.94 | 5.19 | 29,979 | 71.42% | 75.89% |
| RLM-RACE (inner) | miR400 target site | ERR14185198 | 57,898 | 33.01 | 4.81 | 261.01 | 149.87 | 48,929 | 84.51% | 229.69 | 143.24 | 32.9 | 5.14 | 37,637 | 65.01% | 36.44% |
| RLM-RACE (inner) | no target site | ERR14185210 | 79,832 | 29.41 | 4.98 | 235.65 | 113.62 | 68,032 | 85.22% | 195.86 | 103.47 | 28.93 | 5.27 | 41,532 | 52.02% | - |
| RLM-RACE (inner) | no target site | ERR14185214 | 19,856 | 30.45 | 4.87 | 256.86 | 132 | 16,287 | 82.03% | 197.56 | 129.91 | 29.3 | 5.8 | 5,469 | 27.54% | - |
| RLM-RACE (inner) | no target site | ERR14185201 | 10,991 | 30.25 | 4.76 | 279.71 | 155.87 | 8,936 | 81.30% | 221.57 | 154.8 | 29.26 | 5.68 | 1,729 | 15.73% | - |
| RLM-RACE (inner) | no target site | ERR14185197 | 13,253 | 29.97 | 4.68 | 251.49 | 142.84 | 11,167 | 84.26% | 199.99 | 130.11 | 29.4 | 5.29 | 5,781 | 43.62% | - |
| PCR | cDNA PPR1 | ERR14185209 | 75,030 | 29.11 | 3.67 | 569.8 | 89.77 | 363 | 0.48% | 255.17 | 218.55 | 30.52 | 5.54 | 73,878 | 98.46% | - |
| PCR | cDNA PPR1 | ERR14185213 | 63,419 | 29.08 | 3.72 | 545.11 | 108.17 | 344 | 0.54% | 284.39 | 268.94 | 30.78 | 6.38 | 55,778 | 87.95% | - |
| PCR | cDNA PPR1 | ERR14185200 | 58,158 | 28.99 | 3.67 | 558.44 | 117.57 | 374 | 0.64% | 305.33 | 272.26 | 30.32 | 5.83 | 54,313 | 93.39% | - |
| PCR | cDNA PPR1 | ERR14185196 | 47,400 | 29 | 3.7 | 566.32 | 123.13 | 356 | 0.75% | 318.16 | 272.19 | 30.04 | 5.6 | 45,993 | 97.03% | - |
| RLM-RACE (inner) | miR400 target site | ERR14185212 | 72,896 | 34.1 | 4.41 | 285.72 | 115.25 | 63,303 | 86.84% | 250.14 | 104.76 | 34.12 | 4.71 | 56,039 | 76.88% | 54.90% |
| RLM-RACE (inner) | miR400 target site | ERR14185199 | 42,756 | 32.18 | 4.64 | 301.43 | 168.75 | 37,186 | 86.97% | 239.84 | 167.79 | 31.62 | 5.39 | 11,186 | 26.16% | 43.82% |
| RLM-RACE (outer) | Hoxb8 miR-196 target site | ERR14185207 | 12,868 | 27.78 | 4.2 | 281.72 | 84.57 | 1,925 | 14.96% | 255.94 | 148.06 | 27.67 | 4.73 | 864 | 6.71% | 68.14% |
| RLM-RACE (outer) | Hoxa10-AS miR-7162 target site | ERR14185204 | 36,311 | 27.06 | 4.09 | 231.06 | 80.61 | 28,767 | 79.22% | 196 | 62.49 | 26.14 | 4.22 | 25,547 | 70.36% | 76.24% |
| RLM-RACE (inner) | Hoxb8 miR-196 target site A | ERR14185206 | 3,734 | 28.25 | 5.23 | 214.23 | 120.44 | 2,907 | 77.85% | 162.27 | 113.17 | 27.76 | 5.9 | 691 | 18.51% | 15.53% |
| RLM-RACE (inner) | Hoxa10-AS miR-7162 target site A | ERR14185203 | 4,042 | 26.65 | 4.6 | 202.01 | 131.17 | 2,928 | 72.44% | 181.04 | 98.46 | 25.43 | 4.63 | 1,593 | 39.41% | 72.86% |
| RLM-RACE | Hoxa10-AS miR-7162 target site & Hoxb8 miR-196 target site B | ERR14185208 | 57,430 | 28.6 | 4.27 | 215.2 | 95.84 | 39,113 | 68.11% | 255.24 | 104.85 | 27.65 | 4.5 | 31,667 | 55.14% | 73.89% |
| RLM-RACE | Hoxa10-AS miR-7162 target site C | ERR14185205 | 16,708 | 28.15 | 4.5 | 212.94 | 108.71 | 11,441 | 68.48% | 256.29 | 121.58 | 27.34 | 4.97 | 1,693 | 10.13% | 73.84% |
| RLM-RACE-UMI | Hoxa10-AS miR-7162 target site | ERR14172368 | 18,460 | 29 | 3.7 | 234.78 | 176.17 | 8,961 | 48.54% | 148.59 | 121.68 | 28.48 | 5.64 | 2,472 | 13.39% | 45.59% |

Table S2: Cost and throughput of different sequencing platforms. The presented costs are the list prices from the manufacturer's homepages (January 2025).

| Method | Platform | Sequencer cost [€] | Flow cell type | Flow cell cost [€] | Reads per run | Expected Hox8B reads* | Flow cell cost per 1K reads |
| --- | --- | --- | --- | --- | --- | --- | --- |
| PARE | iSeq100 | 20,774 | iSeq 100 i1 Reagent v2 | 643 | 4,000K | 1 | 0.16075 |
| PARE | NextSeq2000 | 326,094 | NextSeq™ 2000 P4 XLEAP-SBS™ Reagent Kit (50 Cycles) | 2,512 | 1,800,000K | 251 | 0.00139556 |
| PARE | NovaSeq X | 984,997 | NovaSeq™ X Series 25B Reagent Kit | 16,000 | 25,000,000K | 3490 | 0.00064 |
| NIL-TDS | MinION Mk1B + Flongle Starter pack | 1,165** | Flongle Flow Cell (R10.4.1) | 66 | 500K | 500,000 | 0.132 |
| NIL-TDS | MinION Mk1B | 570 ** | MinION Flow Cell (R10.4.1) | 665 | 8,000K | 8,000,000 | 0.083 |

*To calculate the expected HoxB8 read depth of a PARE approach, the number of reads per run were multiplied by 2 / 14,323,668, according to Shin et al (2010).

**ONT sells sequencers in a package with library kits and flow cells. For comparability flow cell prices were subtracted from package list prices.

RScript S1. A script to easily evaluate NIL-TDS data and to produce t-plots.

#Example script for "Native Index Ligation-based Targeted Degradome Sequencing (NIL-TDS) for High

Precision Quantification of small RNA Slicing Activity"

#Bernhard T. Werner, Sabrine Nasfi, , M. Lienhard Schmitz, Manar Makhoul, Jens Steinbrenner, Patrick Schäfer

\#

#The script is intended to produce from a basecalled and demultiplexed ONT sequencing run target plots.

#The script utilizes the system2 function to run cutadapt and bowtie2 which need to be installed.

#The script also uses the R-packages ggplot2, ggpubr and wesanderson.

#The options list is used to provide all relevant information to the script

Options<-list()

Options$fastq<-"your_path" #Setting the full path to the fastq file

Options$GOI<-"your_fasta" #Setting the path to your mRNA of interest in fasta

Options$PrimerBS<-"L Position of the inner primer binding site" #Here the binding site of the inner primer should be provided AS INTEGER

#Trimming the reads

i<-Options$fastq

system2("cutadapt",

args=c("-a","TTTCATCAAAGCCAGCAAACGCA",

"--rc",

"-O","20",

"--trimmed-only",

"-o",gsub("\\.fastq","\\.trimmed\\.fastq",i), #Trimmed reads are saved to the same path with the addition trimmed

i

),

stdout=paste0(i,".cut.log"), #Log files are created in the respective folder

stderr = paste0(i,".cut.stderr"),

wait=T)

#Bowtie2 index is generated

system2("bowtie2-build",

args=c("-f",Options$GOI,

Options$GOI))

#Reads are aligned to GOI

i<-gsub("\\.fastq","\\.trimmed\\.fastq",Options$fastq)

system2("bowtie2",

args=c("-q",

"--end-to-end",

"-x",Options$GOI,

"-p","12",

"--mp","6,2",

"--rdg","5,3",

"-k","1",

"--score-min","L,0,-0.6",

"-U",i

),

stdout=gsub("fastq","bt2_out",i), #Pseudo sam is save to the respective folder

stderr=gsub("fastq","bt2.stderr",i), #Log files are created in the respective folder

wait=T

)

#Alignments are read into R

k<-gsub("fastq","bt2_out",i)

#Due to variable header size the first x lines of the sam are skipped until a proper table is generated

sk=5

alignment<-read.delim(k,skip=sk,header=F,quote="",comment.char = "")

while(!ncol(alignment)==19){sk=sk+1

alignment<-read.delim(k,skip=sk,header=F,quote="",comment.char = "")}

names(alignment)<-c("read_name","flag","transcript","position_1","mapq","cigar", "x1","x2","x3","sequence","qual","aln_score","n_ambig_ref",

"n_MM","n_GO","n_GE","Edit_distance","algn_string","f_string")

#Counting of reads for each position

figure_df<-data.frame(reads=integer(),

position=numeric())

for(ii in c((Options$PrimerBS-1100):(Options$PrimerBS-100))){

if(sum(alignment$position_1==(ii))==0){next}

figure_df1<-data.frame(reads=sum(alignment$position_1==(ii)),

position=ii)

figure_df<-rbind(figure_df,figure_df1)

}

#T-plot is generated

library(ggplot2)

p1<-ggplot(figure_df,aes(x=position,y=reads))+

geom_point(alpha=0.5)+

theme_minimal()+

ylab("Reads")+

xlab("Position on GOI")+

theme(panel.border = element_rect(colour = "black", fill=NA, size=1),

plot.margin = margin(0.1,0.1,0.5,0.1, "cm")

)

p1
